# LoFT-TCR: A LoRA-based Fine-tuning Framework for TCR-Antigen Binding Prediction

**DOI:** 10.1101/2025.09.25.678699

**Authors:** Rui Niu, Xiaoying Kong, Xuequn Shang

**Affiliations:** School of Computer Science, Northwestern Polytechnical University, Xi’an, PR China

**Author notes:** Joint first authors (Rui Niu, Xiaoying Kong).

**Keywords:** TCR-antigen binding, Large Language Model, TCR Specificity, Low-Rank Adaptation

## Abstract

T cells recognize and eliminate diseased cells by binding their T cell receptors (TCRs) to short endogenous peptides (antigens) presented on the cell surface. Such interactions are central to adaptive immunity, yet current experimental approaches to identify TCR-antigen binding pairs remain labor-intensive and constrained by limited reagents. Here, we propose LoFT-TCR, a low-rank adaptation (LoRA)-based fine-tuning framework designed for TCR-antigen binding prediction. To capture precise and informative sequence representations, we first fine-tuned the protein large language model ESM-2 on antigen-specific TCR datasets using LoRA. Subsequently, we constructed a heterogeneous interaction graph where nodes encode sequence features and edges indicate interaction relationships. By leveraging a graph learning framework, LoFT-TCR effectively integrates sequence and topological information to enhance prediction capability. Systematic experiments validated that fine-tuning ESM-2 significantly enhanced the model’s capability to extract discriminative sequence representations, which are critical for accurate TCR specificity prediction. Moreover, LoFT-TCR consistently achieved superior performance compared to state-of-the-art methods on both TCR-antigen binding prediction and TCR specificity discrimination tasks. Experimental results demonstrate that LoFT-TCR achieves substantial improvements in predictive accuracy and holds potential for advancing personalized T cell-based immunotherapy.

## I. INTRODUCTION

The antigen-specific recognition by CD8+ T cells of diseased cells plays a key role in the effective eradication of viruses and tumors. T cells utilize T cell receptors (TCRs) to recognize peptides presented by major histocompatibility complex class I (MHC-I) molecules on diseased cells, thereby initiating targeted T cell responses. Peptides capable of activating T cells are known as antigens. Building on the specificity of TCR-antigen recognition, the development of targeted immunotherapies has transformed the landscape of cancer treatment over the past decade [1]. One of the primary methods for identifying TCR-antigen interactions is the use of tetramer-based assays, which are time-consuming, labor-intensive, and even more challenging to implement for less common MHC-I alleles. Advances in this field, along with the evolution of sequencing technologies, have produced extensive immune repertoire data that are now publicly available in various databases. These developments create new opportunities for computational methods to predict TCR-antigen binding, potentially enhancing efficiency and reducing the costs associated with experimental validation.

Focusing on the TCR-antigen binding problem, computational models can be broadly categorized by their learning strategies. Unsupervised models such as TCRdist [2], iSMART [3], and GIANA [4] cluster TCRs based on sequence similarity to infer antigen specificity. Although these models offer advantages such as discovering antigen-specific patterns without prior knowledge, their performance heavily depends on the quality of the input data. The accumulation of high-quality TCR-antigen binding data has promoted the development of supervised models. An early method, NetTCR- 2.0, employs the BLOSUM50 substitution matrix to encode TCR and antigen sequences, and uses separate convolutional neural networks (CNNs) for each sequence type, followed by a classifier [8]. pMTnet incorporates a transfer learning strategy by combining a pre-trained autoencoder (AE) with an LSTM network to generate embeddings for TCR and peptide-MHC-I (pMHC-I) sequences, followed by differential learning to capture binding-related features [9]. Moreover, PanPep integrates meta-learning and neural Turing machines (NTMs) to construct a disentanglement distillation module capable of encoding previously unseen immune sequences [10]. The recently proposed HeteroTCR model uses a pre-trained CNN module to encode immune sequences, formulates TCR-antigen interactions as a graph, and employs a graph neural network (GNN) to extract binding features [11]. These supervised models typically follow a common pipeline: TCR and antigen sequences are first encoded using biological encoding schemes such as BLOSUM50 or amino acid vocabularies, then processed by neural networks to extract informative representations, which are subsequently used to train classifiers that predict binding probabilities. Overall, the emergence of these models has offered new insights into predicting TCR-antigen recognition and has accelerated progress in computational immunology.

However, existing methods face significant challenges due to the immense diversity of the TCR repertoire and off-target recognition issues. High-throughput sequencing reveals that each healthy adult harbors ∼5–6 million TCRs, with less than 1% shared even between siblings [12]. Shallow feature encoders, limited by scarce training data, primarily capture common amino acid motifs and conserved residues, hindering their generalization to novel or rare TCR fragments. Recent advancements in protein large language models (LLMs) offer new opportunities for generating high-quality feature representations. By pre-training on massive protein corpora, LLMs such as ESM-2 are capable of capturing evolutionary and structural patterns in a self-supervised manner [13]. However, representations directly extracted from pre-trained LLMs can be suboptimal for specific downstream tasks, lacking sufficient task-specific adaptation. Fully fine-tuning LLMs for a specific downstream task not only requires substantial computational resources but is also prone to overfitting due to the limited size of the downstream dataset. Furthermore, one antigen can be recognized by multiple TCRs, and one TCR can recognize multiple antigens due to T cell cross-reactivity [14]. This many-to-many interaction naturally forms a heterogeneous bipartite graph, with antigens and TCRs as nodes and their binding events as edges. This biologically informed graph structure complements sequence-level features by providing rich topological context. Additionally, T cell cross-reactivity can cause an off-target issue, where TCRs mistakenly recognize antigens with sequence similarities to intended targets, potentially inducing autoimmune toxicity. Current approaches predict TCR-antigen binding without distinguishing between general antigens and autoimmune-related antigens (arAgs). Enhanced discrimination of TCR recognition specificity is thus urgently needed to achieve precise and personalized immunotherapy.

Inspired by the parameter-efficient low-rank adaptation (LoRA) technique [15], we propose a **Lo**RA-based **f**ine-**t**uning framework for **TCR**-antigen binding prediction, referred to as LoFT-TCR (see Fig. 1). We fine-tuned the ESM-2 model in a self-supervised manner using LoRA, leveraging the adapted many-to-many interaction naturally forms a heterogeneous bipartite graph, with antigens and TCRs as nodes and their binding events as edges. This biologically informed graph structure complements sequence-level features by providing rich topological context. Additionally, T cell cross-reactivity can cause an off-target issue, where TCRs mistakenly recognize antigens with sequence similarities to intended targets, potentially inducing autoimmune toxicity. Current approaches predict TCR-antigen binding without distinguishing between general antigens and autoimmune-related antigens (arAgs). Enhanced discrimination of TCR recognition specificity is thus urgently needed to achieve precise and personalized immunotherapy.

**Fig. 1.**
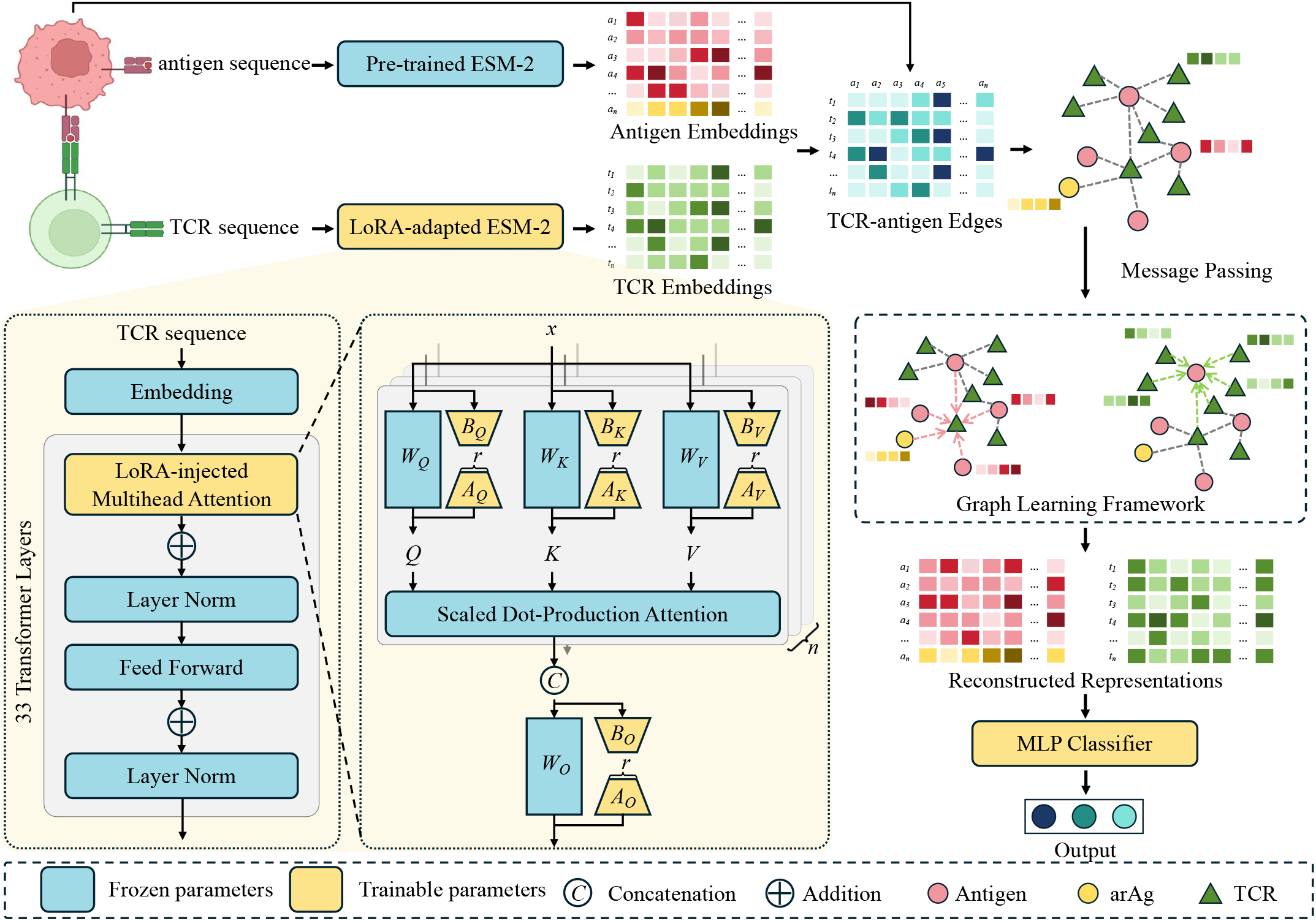
Schematic of the LoFT-TCR framework. Antigen and TCR sequences are encoded by pre-trained or LoRA-adapted ESM-2 models to obtain sequence embeddings. These embeddings are used to construct a TCR–antigen interaction graph, which is further processed by a graph neural network to extract topological features. The combined representations are then fed into an MLP classifier to predict TCR–antigen binding or to classify TCR recognition specificity as general antigens or arAgs.

Inspired by the parameter-efficient low-rank adaptation (LoRA) technique [15], we propose a **Lo**RA-based **f**ine-**t**uning framework for **TCR**-antigen binding prediction, referred to as LoFT-TCR (see Fig. 1). We fine-tuned the ESM-2 model in a self-supervised manner using LoRA, leveraging the adapted model to extract informative sequence representations. Subsequently, we constructed a TCR-antigen interaction graph and applied graph neural networks (GNNs) to extract complementary topological features. These were combined with the sequence representations to train a downstream multi-class multilayer perceptron (MLP) classifier for TCR-antigen binding prediction and TCR recognition specificity discrimination.

The major contributions of our work are three-fold. First, we effectively fine-tuned ESM-2 on an antigen-specific TCR dataset using LoRA, offering a parameter-efficient approach to TCR specificity modeling. Second, we developed LoFT- TCR, a model integrating sequence-level representations from the fine-tuned language model with graph-derived topological features. This integration allows LoFT-TCR to generate comprehensive representations of TCR-antigen pairs. Finally, we demonstrated that LoFT-TCR outperforms existing state-of-the-art methods in both TCR-antigen binding prediction and antigen specificity discrimination tasks.

## II. METHODS

In this section, we present a detailed description of the LoFT-TCR model. As illustrated in Fig. 1, LoFT-TCR consists of three main components. First, the ESM-2 model is fine-tuned on antigen-specific TCRs using LoRA, producing informative sequence representations. Second, we construct a heterogeneous interaction graph using sequence embeddings and known TCR-antigen interactions. A graph learning framework is subsequently employed to complement sequence-level features with topological context. Finally, the integrated representations are fed into a MLP classifier designed for TCR-antigen binding prediction and antigen specificity discrimination tasks.

### A. LoRA-Based Fine-Tuning of ESM-2

The ESM-2 family comprises several protein LLMs pretrained on massive protein corpora using a masked language modeling (MLM) objective. By learning rich, high-dimensional representations of protein sequences, these models enable direct predictions of structural and functional properties from sequence data. Each ESM-2 model is constructed as a stack of *L* transformer layers, with different variants varying in the number of layers and parameters. In this work, we employ the ESM-2 (650M) variant, comprising 33 transformer layers and approximately 650 million parameters, balancing model accuracy and computational efficiency (see Fig. 1).

Formally, for the *l*-th transformer layer, the input is denoted as *x*^(*l*)^ ∈ ℝ^*L×d*^, where *L* is the (padded) sequence length and *d* is the embedding dimension. The queries (*Q*^(*l*)^), keys (*K*^(*l*)^), and values (*V* ^(*l*)^) are computed via linear projections:

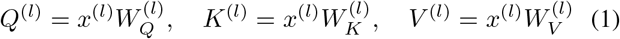

where 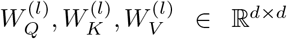 are learnable parameter matrices. For multihead attention with *n* heads, each representation is reshaped into *n* segments of dimension *d*_*k*_, where *d* = *nd*_*k*_, and *d*_*k*_ is the dimensionality per head.

The self-attention output for each head is then calculated as:

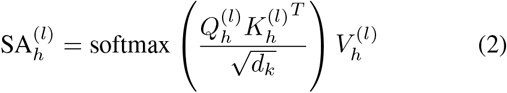

enabling each residue in the sequence to integrate contextual information from all other residues.

ESM-2 employs multihead self-attention, where outputs from all *n* heads, 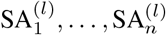, are concatenated and projected back to the original embedding dimension:

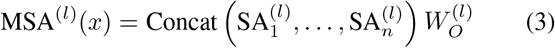

where 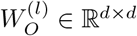 is a learnable output projection matrix.

To efficiently adapt the ESM-2 model to antigen-specific tasks, we utilize the Low-Rank Adaptation (LoRA) technique. To efficiently fine-tune ESM-2, we adopt the LoRA strategy. LoRA freezes all pre-trained weights and introduces additional trainable low-rank matrices specifically into the linear projection layers (queries, keys, values, and output projections) within each transformer block (Fig. 1). This parameter-efficient approach substantially reduces the number of trainable parameters, mitigating overfitting risks particularly in scenarios with limited labeled data.

Formally, for a given pre-trained weight matrix *W*_0_ ∈ ℝ^*d×k*^, LoRA parameterizes the update as a product of two low-rank matrices, *A ∈* ℝ^*r×k*^ and *B ∈* ℝ^*d×r*^, where *r ≪* min(*d, k*):

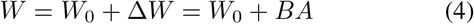

Here, *A* and *B* are the only trainable parameters introduced by LoRA. Typically, matrix *A* is initialized from a Gaussian distribution, while matrix *B* is initialized to zeros. During fine-tuning, only the LoRA parameters *A* and *B* are updated, while the pre-trained weights *W*_0_ remain fixed throughout training. The CDR3 region of the TCR*β* chain is the primary determinant of antigen specificity and diversity, critically influencing antigen recognition efficiency. To accurately capture TCR sequence specificity, we fine-tuned the ESM-2 protein language model, which consists of *L* stacked transformer layers, using an antigen-specific TCR dataset 𝒟. In our implementation, LoRA adapters were integrated into the query (Q), key (K), value (V), and output (O) projection matrices within the self-attention mechanism of each transformer layer. For each targeted module, the LoRA matrices *A* and *B* were initialized using a random Gaussian distribution 𝒩(0, *σ*^2^) and zeros, respectively, while the original weights *W* were frozen. During training, only the LoRA parameters *A* and *B* were updated. In this work, we set the LoRA rank *r* and scaling factor *α* to 4 and 16, respectively. This approach allows the model to effectively learn rich, task-specific representations while preserving pretrained backbone knowledge, substantially reducing the number of trainable parameters. The overall procedure is outlined in Algorithm 1.

Let *f*_*θ*_ denote the original ESM-2 model parameterized by *θ*, mapping an input token sequence *x* ∈ *𝒟* to high-dimensional representations. With the low-rank parameters of LoRA adapters denoted as Δ*θ*, the fine-tuned model is represented as *f*_*θ*+Δ*θ*_, where backbone parameters *θ* remain frozen during fine-tuning.

#### Algorithm 1

**LoRA-based Fine-Tuning of ESM-2**

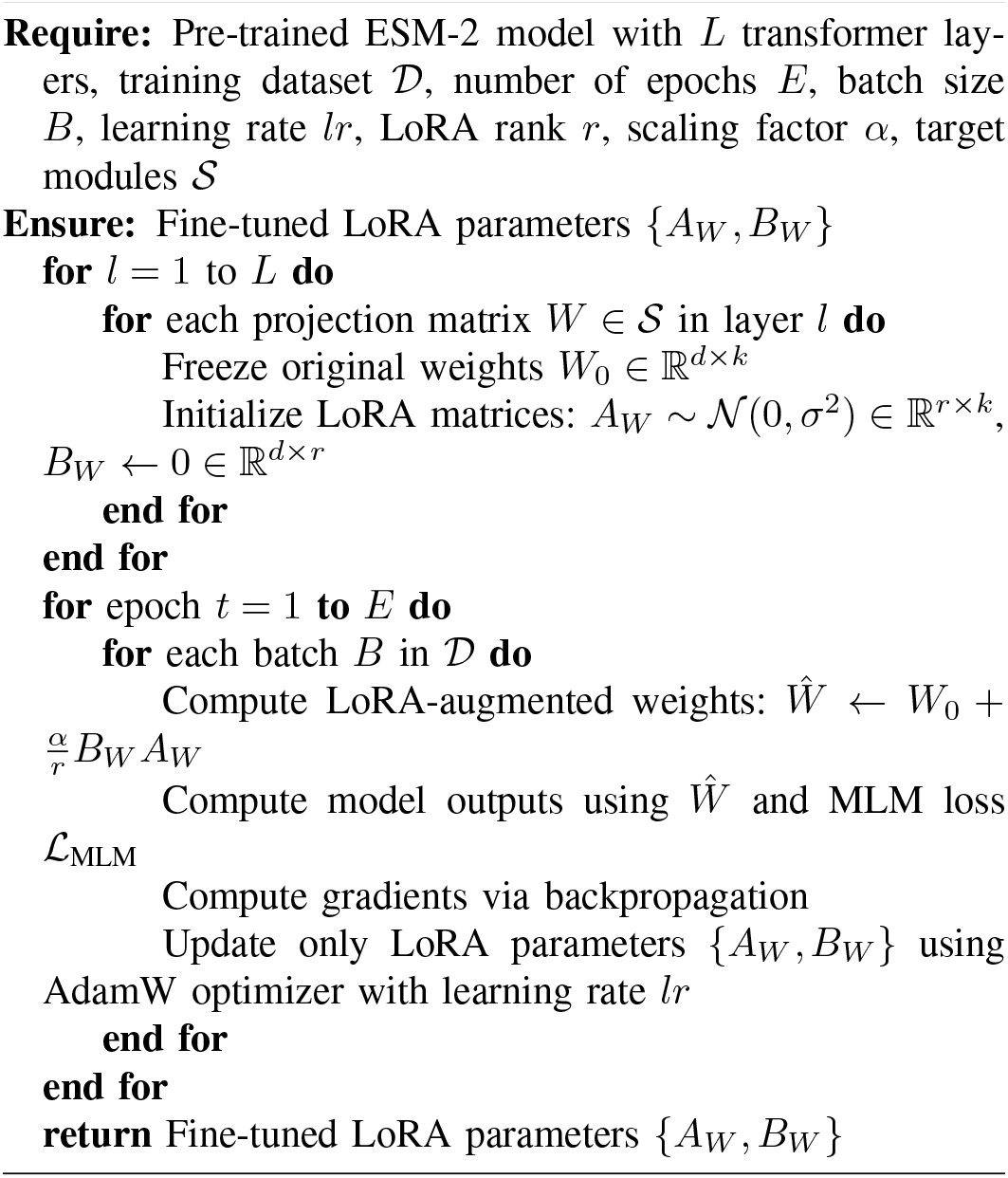

We fine-tuned the model using the MLM objective, computing the MLM loss only over masked token positions as follows:

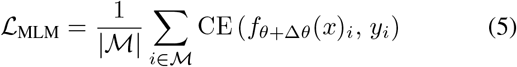

where ℳ represents the set of masked positions, CE denotes the cross-entropy loss, and *y*_*i*_ indicates the true token of position *i*.

### B. Topological Feature Extraction via Graph Learning

To extract topological features from TCR–antigen interactions, we model these interactions as a heterogeneous bipartite graph 𝒢= (𝒩,ℰ). Here, the node set consists of two disjoint subsets: TCR nodes 𝒯 and antigen nodes 𝒜. The edge set ℰ comprises positive edges indicating known binding interactions (*e*^+^), and negative edges representing computationally generated non-binding interactions (*e*^*−*^).

Each node in the graph is represented by a high-dimensional embedding generated by ESM-2. Specifically, for each TCR *t ∈* 𝒯, we employ the fine-tuned model *f*_*θ*+Δ*θ*_ to generate node embedding *h*^0^*t* ∈ ℝ^*d*^. For each antigen *a ∈* 𝒜, the embedding *h*^0^*a ∈* ℝ^*d*^ is extracted using the original pre-trained ESM-2 model *f*_*θ*_. This design integrates both task-specific and general protein representations into the graph learning framework.

We employed a GNN framework on this heterogeneous graph for iterative message passing and representation learning. Specifically, node embeddings are iteratively updated by aggregating information from their immediate neighborhoods. For each antigen *a ∈* 𝒜, the embedding at layer *l* + 1 is computed by:

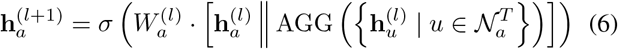

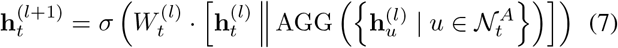

where 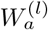 and 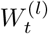 are trainable weight matrices, *σ*(*·*) indicates sigmoid activation function, AGG(*·*) denotes mean aggregation over the neighborhood embeddings, 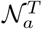 is the set of TCR neighbors of antigen *a*, and 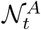 is the set of antigen neighbors of TCR *t*. This design enables the model to jointly capture antigen-specific recognition patterns in TCRs and TCR-binding motifs in antigens, leveraging both sequence-derived and topological information.

### C. TCR-antigen Specificity Prediction

After graph propagation, the informative representations of TCR and antigen were concatenated and fed into a downstream MLP classifier. Specifically, we implemented two versions of the MLP classifier: one for TCR–antigen binding prediction and another for classifying TCR recognition specificity into general antigens and arAgs. Both classifier branches share the same backbone architecture, which consists of two fully connected layers with ReLU activations; the first layer contains 512 neurons and the second contains 256 neurons.

The output layers of the two classifiers were designed differently. For the TCR–antigen binding prediction task, the output layer consists of a single neuron with sigmoid activation, where the output probability 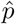 indicates the likelihood of binding for the given TCR–antigen pair. For the TCR recognition specificity classification task, the labels are defined as *p ∈* {0, 1, 2}, representing non-binding, recognition of general antigens, and recognition of arAgs, respectively. Accordingly, the output layer contains three neurons with softmax activation, producing a probability vector 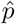 over the three classes.

Finally, the cross-entropy loss function was used for both the binary (binding prediction) and multi-class (recognition specificity) classification tasks.

## III. EXPERIMENTS

### A. Dataset

To facilitate the fine-tuning of ESM-2 for capturing TCR specificity, we curated a benchmark dataset by aggregating TCR-antigen binding records from three widely used public databases:

- **IEDB**, which archives a broad spectrum of experimentally validated T cell epitopes associated with infectious diseases, allergies, autoimmune disorders, and cancers [5];
- **McPAS-TCR**, a manually curated collection of TCR sequences linked to diverse antigenic targets across various pathologies [6];
- **VDJdb**, a curated repository of TCRs annotated with known antigen specificities, accompanied by confidence scores to quantify annotation reliability [7].

We restricted our dataset to human-derived TCR*β* CDR3 sequences associated with human leukocyte antigen class I (HLA-I) molecules. Filtering was performed to ensure data quality and biological relevance. We retained TCR*β* CDR3 sequences with lengths between 10 and 20 amino acids and included only peptides ranging from 8 to 15 residues. Entries containing lowercase characters, non-standard amino acids, or duplicated TCR-antigen pairs were excluded. To improve antigen-level specificity, we applied iSMART to cluster TCR*β* sequences based on pairwise local alignment and excluded singletons that did not associate with any cluster. After preprocessing, the resulting dataset contained 41,732 high-confidence TCR-antigen binding pairs, including 972 distinct peptides and 37,054 unique TCR*β* CDR3 sequences.

We further curated arAgs from two publicly available databases: IEDB and McPAS-TCR. From IEDB, we selected peptides associated with autoimmune conditions in human subjects and restricted to HLA-I alleles. From McPAS-TCR, we retrieved peptides annotated under the “autoimmune” disease category and derived from human samples. The collected peptides were then merged and processed using the same filtering criteria as described for the benchmark dataset. This resulted in a total of 786 curated arAgs.

### B. Inplementation Details

All experiments were conducted on NVIDIA A800-N GPUs, each equipped with 80GB of GPU memory.

We fine-tuned the ESM-2 (650M) ^1^ model using LoRA-based adaptation within the HuggingFace Transformers framework, accelerated by DeepSpeed Stage-2 ZeRO optimization and FP16 mixed-precision training. AdamW optimization was used, configured with a learning rate of 2e-5, batch size of 32, and trained for one epoch.

LoFT-TCR was also trained using FP16 mixed-precision. The model was trained for 1,000 epochs using full-batch gradient descent optimized by Adam at a learning rate of 1e-3. To guarantee fair comparisons and reproducibility, baseline models were strictly implemented according to their official repositories.

### C. learning TCR Specificity

To enhance the representational capacity of the pre-trained ESM-2 model for capturing detailed TCR specificity, we fine-tuned it on 37,054 antigen-specific TCR sequences using the LoRA adapter. Specifically, we randomly allocated 90% of the TCR sequences to the training set and the remaining 10% to the validation set. To mine TCR specificity in a self-supervised manner, we fine-tuned ESM-2 using the MLM objective, randomly masking 15% of tokens and predicting these masked tokens. To prevent potential overfitting, we limited the fine-tuning to one epoch. The accuracy for predicting masked tokens is calculated as follows:

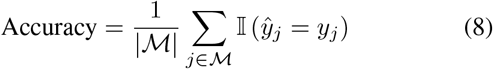

where ℳ is the set of masked token indices, 𝕀(*·*) is the indicator function, which evaluates to 1 if its argument is true and 0 otherwise. The 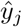 and *y*_*j*_ are predicted and true label for position *j*, respectively.

The training and validation losses and accuracies are illustrated in Fig. 2, showing clear convergence trends during finetuning. We observed that the training loss decreased slowly during the initial 40 steps, dropped rapidly between steps 40 and 100, indicating rapid feature learning, and plateaued after 100 steps. Correspondingly, accuracy remained stable during the first 40 steps, improved rapidly between steps 40 and 100, and then gradually increased.

**Fig. 2.**
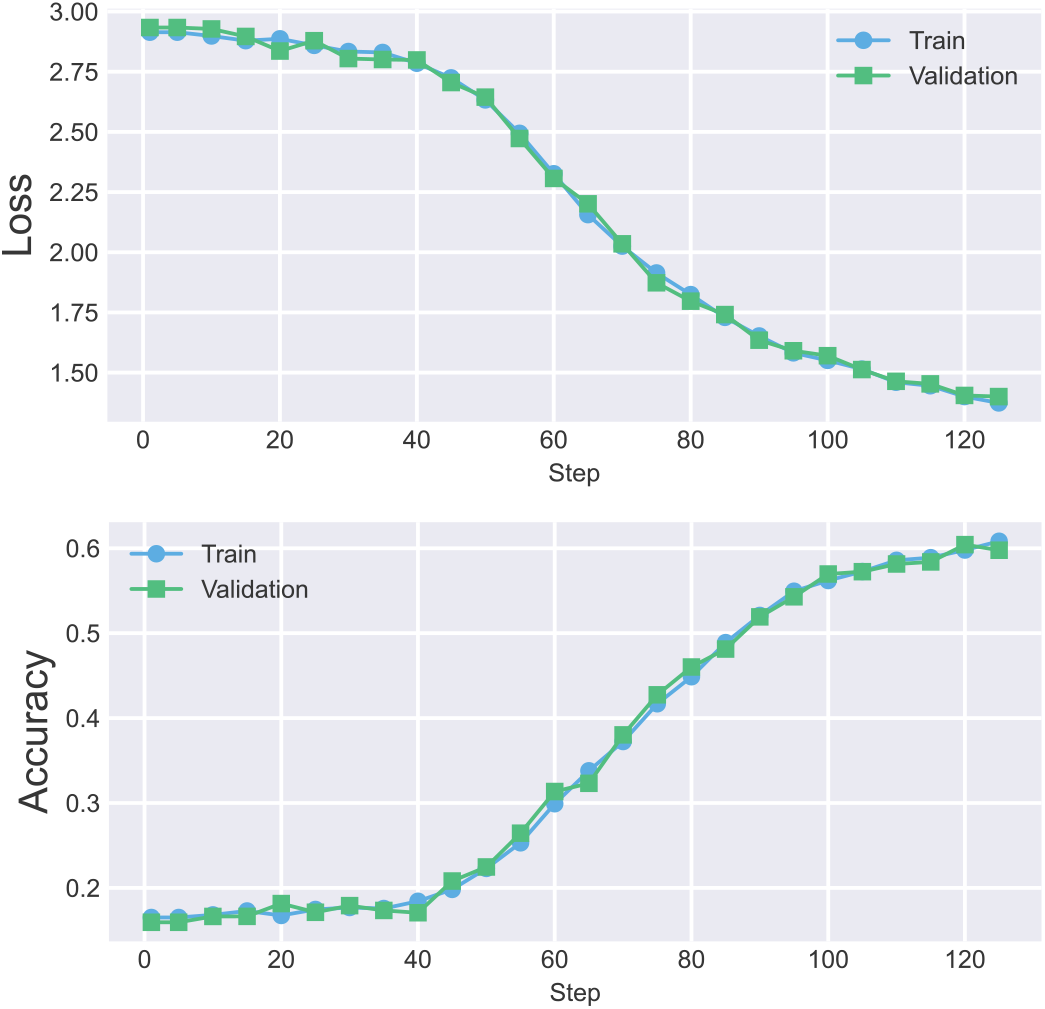
Training and validation loss and accuracy curves for the LoRA-adapted ESM-2 model.

**Fig. 3.**
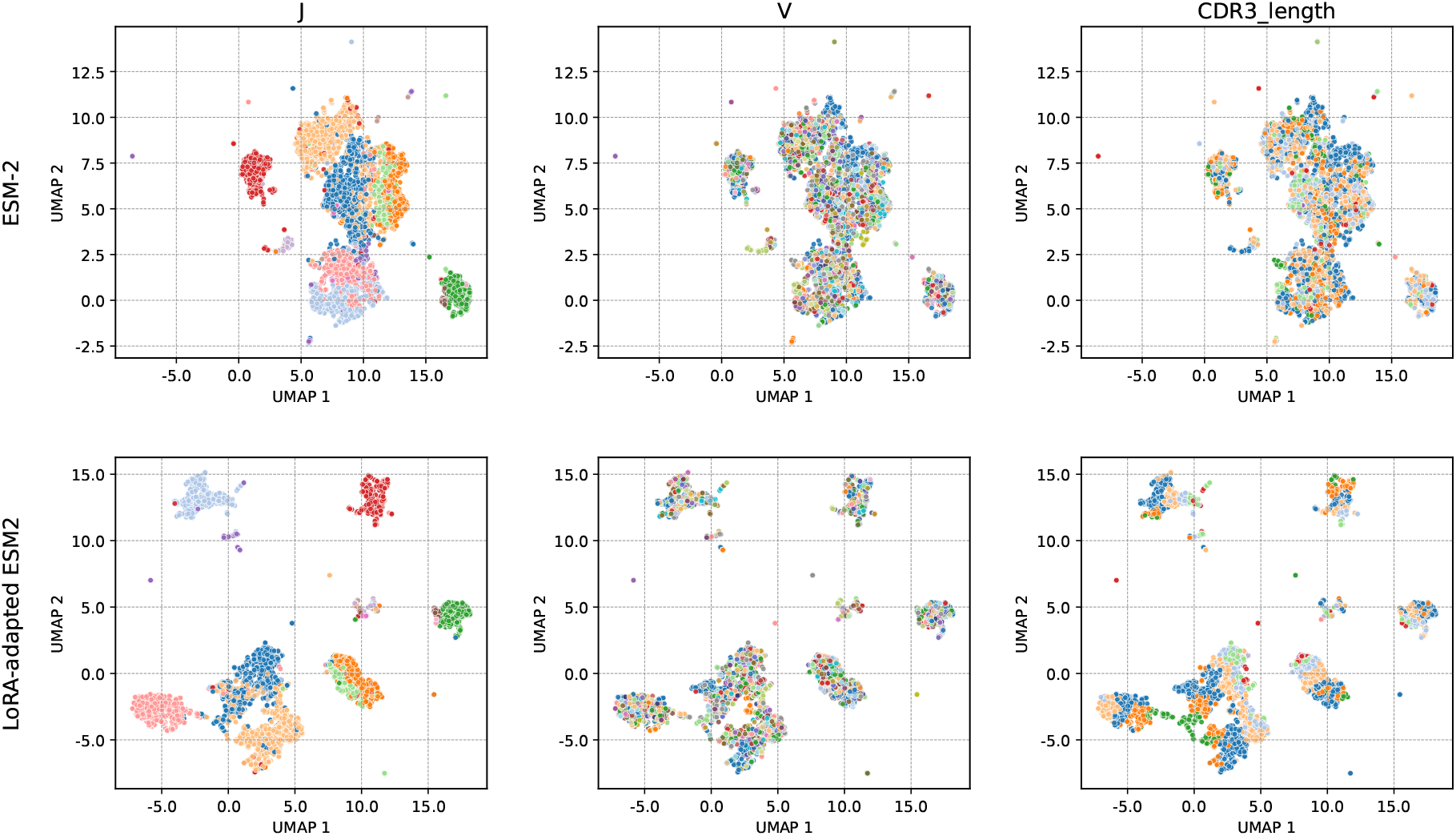
UMAP projections of TCR embeddings obtained from the pre-trained and LoRA-adapted ESM-2 models. Each point denotes a TCR from the VDJdb dataset and is colored by J gene usage, V gene usage, or CDR3 length in separate panels.

The LoRA adapter fine-tunes ESM-2 by injecting low-rank matrices into specific target modules within the multi-head attention mechanism. To identify optimal target modules for efficient fine-tuning, we conducted an ablation study by progressively reducing the target modules (Q, K, V, O) and comparing computational costs and model performance. We also evaluated the original ESM-2 model without fine-tuning as a baseline (Table I). All three fine-tuned models outperformed the original model on the validation set after one epoch. Specifically, the Q-only targeted model reduced the trainable parameters by 88.9% compared to the full-module targeted model but provided minimal accuracy improvement over the baseline. The Q/V-targeted model reduced parameters by 77.8%, yet exhibited a 29.1% decrease in accuracy compared to the fully targeted model. Extending training epochs, we observed that the Q-targeted model reached an accuracy of 0.607 after two epochs, while the Q/V-targeted model plateaued at 0.548 after three epochs. Consequently, we selected the fully targeted version (Q, K, V, O) for subsequent analyses.

**TABLE I.** Comparison of targeting different combinations of multihead attention modules in LoRA fine-tuning.

| Metric | Weight Type |  |  |  |
| --- | --- | --- | --- | --- |
| | Baseline | $W_Q(\uparrow)$ | $W_Q, W_V(\uparrow)$ | $W_Q, W_K, W_V, W_O(\uparrow)$ |
| Number of trainable parameters | 0 | 337,920 | 675,840 | 3,051,520 |
| Accuracy | 0.176 | 0.180 | 0.307 | **0.598** |
| DB index (V gene) | 67.3 | 46.8 | **46.7** | 58.7 |
| DB index (J gene) | 3.93 | 2.59 | 2.17 | **1.86** |
| DB index (Length) | 35.2 | 10.5 | 24.24 | **10.4** |

Note: ( ↑ ) indicates that using the corresponding weight type leads to better performance compared to the baseline.

We next employed a dataset containing detailed annotations of TCR sequences to evaluate whether fine-tuning could enhance the model’s ability to capture TCR specificity. We collected antigen-specific TCR sequences from VDJdb [7], retaining only those with a confidence score greater than 0 and sequence lengths between 10 and 20 amino acids. The resulting dataset comprised 9,971 unique TCR sequences annotated with 14 distinct V genes, 53 distinct J genes, and corresponding sequence lengths.

The extensive diversity of the TCR repertoire primarily arises from the recombination of V(D)J genes during T-cell development. V genes encode variable regions that form the primary domains responsible for antigen recognition by TCRs. J genes encode joining regions that similarly contribute to generating TCR diversity. Additionally, TCR sequence length influences the structural conformation of the receptor, subsequently affecting its antigen recognition capability. To quantitatively assess the effectiveness of fine-tuning, we introduced the Davies–Bouldin (DB) index, which measures cluster separability by evaluating intra-cluster compactness and intercluster separation in the embedding space.

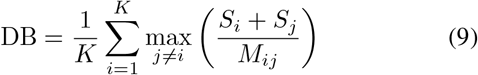

As shown in Table I, all three fine-tuned models exhibited lower DB index values (indicating better cluster separability) for V gene usage, J gene usage, and sequence length compared to the baseline (unfine-tuned) model. This result demonstrates that fine-tuning via self-supervised training effectively captures TCR specificity information.

After fixing all the parameters, we fine-tuned the ESM-2 model using all antigen-specific TCR sequences as the training dataset. The resulting model was used in subsequent experiments.

### D. TCR-antigen Binding

First, we evaluated the performance on the general TCR-antigen binding prediction task without discriminating between general antigens and arAgs. We employed 5-fold cross-validation for a fair comparison of LoFT-TCR with other baseline models. The dataset comprising 41,732 experimentally validated TCR-antigen binding pairs was used as positive samples and equally partitioned into five folds. Due to the scarcity of experimentally validated non-binding TCR-antigen pairs in existing databases, negative samples were artificially generated in each fold. Specifically, for each antigen, we constructed an equal number of negative samples by randomly pairing the antigen with TCR sequences not known to interact with it.

As illustrated in Fig. 1, LoFT-TCR leverages LoRA-adapted ESM-2 as the feature encoder and incorporates graph learning to integrate topological information into the representations. To systematically evaluate the effectiveness of our model architecture, we compared LoFT-TCR against several representative methods employing diverse feature encoders and graph learning strategies, including NetTCR-2.0 [8], pMTnet [9], and HeteroTCR [11]. NetTCR-2.0 employs a BLOSUM50- based encoding of antigen and TCR sequences, subsequently utilizing a shallow CNN for feature extraction and prediction. Although NetTCR-2.0 accepts both TCR *α* and *β* chains as input, we restricted our analysis to the TCR *β* chain, which predominantly determines antigen recognition specificity. Following a transfer-learning strategy, pMTnet initially pre-trained an autoencoder on an external TCR dataset and a Long Short-Term Memory (LSTM) network on an external pMHC-I dataset. These pre-trained feature encoders were integrated into a unified model, whose output embeddings were fed into a fully connected neural network to predict TCR-antigen recognition. In our experiments, we directly utilized these pre-trained encoders without further fine-tuning. Although the LSTM encoder was originally trained on the pMHC-I dataset, we adopted it as the antigen feature encoder. The recently proposed HeteroTCR model employs two CNNs, pre-trained separately on TCR and antigen sequences, as feature encoders. It also integrates a graph learning module to capture topological relationships.

To further assess the respective contributions of the LLM-based feature encoder and the graph learning framework, we conducted an ablation study involving two variants of LoFT- TCR. In the “GNN + MLP” variant, we replaced the ESM- 2 encoder with the biologically informed encoding scheme BLOSUM50. In the “ESM-2 + MLP” variant, we removed the GNN module, directly feeding feature embeddings from ESM-2 into the MLP classifier.

We employed area under the receiver operating characteristic curve (AUROC), area under the precision-recall curve (AUPRC), and accuracy metrics to systematically evaluate model performance. As shown in Table II, LoFT-TCR consistently outperformed all baseline models across all evaluation metrics. Compared with HeteroTCR, which also employs a graph learning framework combined with pre-trained feature encoders, LoFT-TCR still achieved improvements of 6.6% and 7.6% in average AUROC and AUPRC, respectively, highlighting the effectiveness of the fine-tuned ESM-2 encoder. The ablation study demonstrated that both high-quality sequence representations and topological information significantly contributed to LoFT-TCR’s performance. Specifically, removing the ESM-2 encoder or the GNN module led to performance declines of 13.8% and 10.3% in AUROC, respectively. These results effectively demonstrate the robustness and superior predictive capability of LoFT-TCR in the TCR-antigen recognition task.

**TABLE II.** Performance comparison on TCR–antigen binding and specificity prediction.

| LLM Module | Feature Encoder | Graph-Based Modeling | Methods | TCR–antigen binding prediction |  |  | TCR–antigen specificity discrimination |  |  |
| --- | --- | --- | --- | --- | --- | --- | --- | --- | --- |
|  |  |  |  | AUROC | AUPRC | Accuracy | macro AUROC | macro AUPRC | macro Accuracy |
| – | ✓ | – | NetTCR-2.0 | 0.758±0.004 | 0.763±0.004 | 0.680±0.004 | – | – | – |
| – | ✓ | – | pMTnet | 0.615±0.012 | 0.602±0.011 | 0.580±0.007 | – | – | – |
| – | ✓ | ✓ | HeteroTCR | 0.793±0.006 | 0.801±0.007 | 0.709±0.004 | – | – | – |
| – | – | ✓ | GNN+MLP | 0.721±0.009 | 0.752±0.007 | 0.622±0.004 | 0.818±0.006 | 0.697±0.010 | 0.643±0.005 |
| ✓ | ✓ | – | ESM-2+MLP | 0.756±0.007 | 0.758±0.008 | 0.672±0.003 | 0.849±0.003 | 0.761±0.003 | 0.670±0.004 |
| ✓ | ✓ | ✓ | **LoFT-TCR** | **0.859±0.002** | **0.877±0.002** | **0.758±0.004** | **0.912±0.004** | **0.869±0.007** | **0.757±0.008** |

Note: “✓” indicates that the model includes the corresponding component. Performance values are reported as mean ± standard deviation across 5-fold cross-validation. The best experimental results are marked with **Bold Font**.

### E. TCR-antigen Specificity Discrimination

We then evaluated LoFT-TCR’s capability to distinguish between TCR recognition specificities towards general antigens and arAgs. Current approaches typically focus on predicting TCR-antigen binding without differentiating between general antigens and arAgs. However, due to T-cell cross-reactivity, TCRs may recognize arAgs sharing sequence similarities with the intended antigens, leading to a significant off-target issue. Therefore, developing a method to accurately discriminate TCR specificity between intended antigens and arAgs is highly desirable.

We labeled the 41,732 experimentally validated TCR-antigen pairs as either general antigen-binding or arAg-binding according to their recognition specificities. We employed 5- fold cross-validation for performance evaluation of LoFT- TCR based on this labeled dataset. Negative samples were artificially generated following the same procedure used in the binding prediction task.

To simultaneously predict TCR binding to general antigens, arAgs, and non-binding interactions, we applied a trinary version of LoFT-TCR, in which the output layer consists of three neurons. As no existing method explicitly addresses TCR specificity discrimination, we selected two previously established variants for comparison. We modified their output layers accordingly to accommodate the three-class prediction task.

We utilized macro AUROC, macro AUPRC, and macro accuracy metrics to comprehensively evaluate the models’ performance in the multi-class prediction task. As shown in Table II, LoFT-TCR achieved average macro AUROC, AUPRC, and accuracy scores of 0.912, 0.869, and 0.757, respectively, clearly outperforming the two comparative variants. We observed that the feature-based variant achieved improvements of 3.1% and 6.4% in macro AUROC and AUPRC, respectively, over the graph-based variant. This result aligns with our findings from the TCR-antigen binding task, further emphasizing that high-quality sequence representation is crucial for improving model predictive capability. These results effectively demonstrate the capability and utility of LoFT-TCR for precise discrimination of TCR recognition specificity between general antigens and arAgs.

## IV. CONCLUSION

In this work, we presented LoFT-TCR, a LoRA-based fine-tuning framework designed to enhance TCR-antigen binding prediction through efficient adaptation of large protein language models. By fine-tuning ESM-2 on antigen-specific TCR datasets, we demonstrated that the model could effectively capture more discriminative sequence representations critical for TCR specificity prediction. LoFT-TCR consistently out- performed state-of-the-art methods on both the TCR-antigen binding prediction and TCR specificity discrimination tasks, validating its effectiveness and superiority. Future research will incorporate additional antigen-specific TCR information, such as *α* chain sequences and V(D)J gene usage, to further enhance the representation capability of the fine-tuned language models. Overall, we believe that LoFT-TCR will contribute to the development of personalized T cell-based immunotherapy by enabling more precise and reliable TCR specificity predictions.

## Acknowledgment

This study was supported by the National Natural Science Foundation of China (Grants No. 62433016 and 624B2112, awarded to X.S., R.N., respectively). We gratefully acknowledge these funding agencies for their support.

## Footnotes

1 https://dl.fbaipublicfiles.com/fair-esm/models/esm2t33650MUR50D.pt

## Notes

### Competing Interest Statement

The authors have declared no competing interest.

